# *NEK1* haploinsufficiency worsens DNA damage but not defective ciliogenesis in *C9ORF72* patient-derived iPSC-motoneurons

**DOI:** 10.1101/2024.03.07.582752

**Authors:** Serena Santangelo, Sabrina Invernizzi, Marta Nice Sorce, Valeria Casiraghi, Silvia Peverelli, Alberto Brusati, Claudia Colombrita, Nicola Ticozzi, Vincenzo Silani, Patrizia Bossolasco, Antonia Ratti

## Abstract

A hexanucleotide G_4_C_2_ repeat expansion (HRE) in *C9ORF72* gene is the major cause of amyotrophic lateral sclerosis (ALS) and frontotemporal dementia (FTD), leading to both loss- and gain-of-function pathomechanisms. The wide clinical heterogeneity among *C9ORF72* patients suggests potential modifying genetic factors. Notably, *C9ORF72* mutations often co-occur with other variants in ALS/FTD-associated genes, such as *NEK1*, which encodes for a kinase involved in multiple pathways including DNA damage response and ciliogenesis. In this study, we generated induced pluripotent stem cells (iPSCs) and differentiated motoneurons (iPSC-MNs) derived from an ALS patient carrying both *C9ORF72* HRE and a *NEK1* loss-of-function mutation to study the effect of the *NEK1* haploinsufficiency on *C9ORF72* pathology. Double mutant *C9ORF72*/*NEK1* cells showed increased pathological *C9ORF72* RNA foci in iPSCs and higher DNA damage levels in iPSC-MNs compared to single mutant *C9ORF72* cells. In contrast, ciliogenesis was similarly impaired in both *C9ORF72* and *C9ORF72/NEK1* iPSC-MNs showing shorter cilia. Altogether, our study supports the use of patient-derived iPSCs to functionally explore the contribution of genetic modifiers in *C9ORF72*-associated pathology.

## 1. Introduction

The hexanucleotide G_4_C_2_ repeat expansion (HRE) in the first intron of *C9ORF72* gene represents the major genetic cause of both amyotrophic lateral sclerosis (ALS) and frontotemporal dementia (FTD) (DeJesus-Hernandez et al., 2011; Renton et al., 2011). *C9ORF72* HRE-associated pathomechanisms comprise both loss- and gain-of-function conditions, the latter characterized by the formation of toxic HRE-containing RNA foci and the translation of dipeptide-repeat proteins (DPRs) which altogether lead to alterations of several cellular pathways (Balendra and Isaacs, 2018). The wide clinical heterogeneity observed among *C9ORF72* patients, encompassing disease manifestation (ALS/FTD), age of onset, progression rate and clinical symptoms, suggests the potential influence of both genetic and epigenetic modifier factors. Among the genetic modifiers, the oligogenicity condition in *C9ORF72* carriers has already been described, although its correlation with specific clinical features still remains uncertain (Ross et al., 2020; Ruf et al., 2023; Van Daele et al., 2023). Of note, *C9ORF72* HRE is frequently found in combination with rare variants in other ALS and/or FTD-associated genes, including *TARDBP, SOD1, FUS* and other minor genes, such as *NEK1* (Nguyen et al., 2018a; Riva et al., 2022). *NEK1* heterozygous loss-of-function (LOF) variants and the missense p.Arg261His variant have been identified in about 3% of both familial and sporadic ALS cases (Brenner et al., 2016; Kenna et al., 2016). *NEK1* encodes for a serine/threonine tyrosine kinase involved in the maintenance of genomic stability and DNA damage response (DDR) (Chen et al., 2011), cell cycle regulation (Chen et al., 2008; Pelegrini et al., 2010), mitochondrial activity (Martins et al., 2021) and ciliogenesis (Shalom et al., 2008; White and Quarmby, 2008), although the biological effects of ALS-related *NEK1* mutations on these pathways remain largely unexplored.

Different studies have already used induced-pluripotent stem cells (iPSCs) and differentiated motoneurons (iPSC-MNs) to investigate the effect of *C9ORF72* HRE (Lopez-Gonzalez et al., 2016) and *NEK1* LOF mutations on DNA damage and DDR (Higelin et al., 2018) and on nucleo-cytoplasmic transport (Mann et al., 2023). However, the combined effect of distinct ALS gene variants has been poorly investigated functionally so far and patient-derived iPSC-MNs represent a suitable disease model for this purpose.

## 2. Materials and Methods

### 2.1. iPSC reprogramming and culturing

Primary fibroblasts of the double mutant *C9ORF72*/*NEK1* patient were isolated from skin biopsy as previously described (Onesto et al., 2016) after written informed consent and approval by the ethics committee (2015-03-31-07). iPSC reprogramming was performed using the CytoTune®-iPS 2.0 Sendai Reprogramming Kit (*Thermo Fisher Scientific*) as previously described (Bardelli et al., 2020). The obtained iPSC clones were cultured in E8 medium (*Thermo Fisher Scientific*) and passaged twice a week using 0.5 mM PBS/EDTA. Spontaneous differentiation assay was performed as previously described (Santangelo et al., 2023). Briefly, iPSCs were grown in suspension for 7 days to generate embryoid bodies (EBs) and they were then plated on Matrigel-coated coverslips and grown in E8 medium for additional 10 days. Expression of stemness markers was assessed by reverse-transcription PCR (RT-PCR) using the primers already described (Bardelli et al., 2020). iPSC lines derived from a wild-type healthy individual (Bossolasco et al., 2018) and a mutant *C9ORF72* patient were previously generated and characterized as described (Bardelli et al., 2020).

### 2.2. Genetic analyses

Genomic DNA was extracted from fibroblasts and iPSCs using the Wizard® Genomic DNA Purification Kit (*Promega*). *C9ORF72* HRE was tested by repeat-primed PCR (RP-PCR) as previously described (Ratti et al., 2022). To determine HRE size, Southern Blot analysis was performed using a ^32^P-radioactive probe recognizing a unique region flanking the *C9ORF72* repeat sequence, as previously reported (Ratti et al., 2022). Heterozygous *NEK1* nonsense mutation was assessed by Sanger sequencing using specific primers (FOR_TATTTTCCTGATATGTGGGTTTT and REV_TGGATGTGTGTTTGTGTCTGT) and the amplicon was sequenced on an ABI 3500 Genetic Analyzer (*Applied Biosystems*). Q-banding was used for cytogenetic analysis of the iPSC line as already described (Santangelo et al., 2023).

### 2.3. iPSC-MNs differentiation and DNA damage induction

iPSCs were grown in suspension for 21 days to generate EBs, that were then dissociated and cultured for additional 30 days to obtain iPSC-MNs, as previously described (Bardelli et al., 2020). DNA damage was induced with 20 μM Neocarzinostatin (NCS) (*Merck*) for 20 minutes (Francia et al., 2016) with a subsequent 6-hours rescue timeframe from NCS treatment.

### 2.4. Western Blot

Cell pellets were incubated in lysis buffer (20 mM Tris-HCl pH 7.5, 150 mM NaCl, 1 mM EDTA, 1 mM EGTA, 1% Triton X-100, protease inhibitor cocktail (*Roche*)), sonicated and centrifuged at 13’000 x g at 4 °C for 15 minutes. Protein extracts were quantified using the Pierce™ BCA Protein Assay Kit (*Thermo Fisher Scientific*) and 40 μg of protein lysates were resolved on precast 3-8% gels in Tris-acetate buffer and then transferred to nitrocellulose membranes (all from *Thermo Fisher Scientific*). Membranes were blocked with 5% (w/v) non-fat dry milk in Tris-buffered saline (*Santa Cruz Biotechnology*) with Tween-20 (*Sigma-Aldrich*) (TBST) buffer and incubated with primary antibodies **(Supplementary Table S1)** in blocking solution at 4 °C overnight and with HRP-conjugated secondary antibodies **(Supplementary Table S1)** for 1 hour at room temperature. The Clarity™ Western ECL Substrate (*Biorad*) was used for signal detection and densitometric analyses were performed using the Quantity One software (*Biorad*).

### 2.5. Immunofluorescence

Immunofluorescence (IF) was performed as previously described (Ratti et al., 2022). Briefly, cells were fixed in 4% paraformaldehyde (*Santa Cruz Biotechnology*) for 20 minutes and permeabilized with ice-cold methanol and with 0.3% Triton X-100 for 5 minutes each. Blocking was performed in 10% Normal Goat Serum (*Gibco*) in PBS for 20 minutes at room temperature and cells were incubated with primary antibodies **(Supplementary Table S1)** at 37 °C for 1.5 hours and then with secondary antibodies **(Supplementary Table S1)** for 45 minutes in blocking solution at room temperature. Nuclei were stained with 4’6-diamidino-2-phenylindole (DAPI) (*Merck*).

### 2.6. RNA-Fluorescence in Situ Hybridization

RNA-Fluorescence In Situ Hybridization (RNA-FISH) was performed as previously described (Bardelli et al., 2020). Briefly, fixed cells in 4% paraformaldehyde were permeabilized with 0.2% Triton X-100 for 10 minutes and dehydrated with 70%, 90%, 100% ethanol for 5 minutes each in a desiccant chamber. Pre-hybridization was performed in 50% formamide (*IBI Scientific*), 50 mM sodium phosphate, 10% dextran sulphate (*Merck*) and 2X saline-sodium citrate (SSC) at 66 °C for 1 hour. Hybridization was performed at 66 °C overnight with 40 nM 5′ TYE-563-labelled locked nucleic acid (LNA)-(C_4_G_2_)_2.5_ probe (*Exiqon Qiagen*). Cells were then washed once in 2X SSC/0.1% Tween-20 for 5 minutes and three times in 0.1X SSC for 10 minutes at RT before being dehydrated as above and nuclei stained with DAPI.

### 2.7. Image acquisition and analyses

Images were acquired using the confocal Eclipse Ti microscope (*Nikon*) as Z-stacks (0.5 μm step size for IF images, 0.2 μm step size for RNA-FISH images) at 60X magnification. Images were analysed with the *ImageJ* software (https://imagej.nih.gov/ij). In particular, for quantitative analysis of *C9ORF72* RNA foci and γH2AX foci, the Find Maxima function was used after setting an appropriate threshold, while quantification of primary cilia length was performed using the CiliaQ plugin (Hansen et al., 2021).

### 2.8. Statistical analysis

Statistical analyses were conducted using the Graphpad Prism 9 software, using Student’s t-test, one-way ANOVA or two-way ANOVA with Tukey’s multiple comparison *post hoc* test. Results were considered statistically significant if p≤0.05. Bar charts represent the mean ± standard error mean (SEM).

## 3. Results

### 3.1. Generation and characterization of iPSCs from the double mutant *C9ORF72*/*NEK1* ALS patient

By a genetic screening of a cohort of Italian ALS patients we identified a patient carrying *C9ORF72* HRE and a concurrent heterozygous pathogenetic non-sense mutation in *NEK1* gene (NEK1_NM_001199397: c.3107C>G: p.Ser1036Ter), already demonstrated to cause *NEK1* haploinsufficiency (Nguyen et al., 2018). The clinical features of the ALS *C9ORF72/NEK1* patient are reported in **Supplementary Table S2**.

To investigate the impact of *NEK1* haploinsufficiency as a possible modifying factor of *C9ORF72* pathology at biological level, we generated an *in vitro* disease model by reprogramming the primary fibroblasts of the double mutant *C9ORF72*/*NEK1* patient into iPSCs. The newly generated *C9ORF72*/*NEK1* iPSC line presented a canonical morphology **(Fig. 1a)** and fulfilled stemness features by expressing the typical pluripotency markers SSEA-4, alkaline phosphatase (AP) and TRA-1-60 by IF **(Fig. 1b)** and *SOX2, OCT3/4* and *NANOG* by RT-PCR **(Supplementary Fig. S1a)**. We further assessed the pluripotency of the *C9ORF72*/*NEK1* iPSC line by evaluating its ability to spontaneously differentiate into the three germ layers, characterized by the expression of alpha-fetoprotein (endoderm), βIII-tubulin (ectodem) and desmin (mesoderm) markers **(Supplementary Fig. S1b)**. The ALS double mutant iPSCs had a normal karyotype as assessed by Q-banding **(Fig. 1c)** and genetic analysis confirmed the presence of the HRE in *C9ORF72* gene by RP-PCR **(Fig. 1d)** and of the heterozygous p.Ser1036Ter mutation in *NEK1* gene by Sanger sequencing **(Fig. 1f)**. Southern blot analysis revealed HRE lengths of ∼600 and ∼200 units in the original patient’s fibroblasts, indicating the presence of somatic mosaicism, and of ∼500 units in the reprogrammed iPSC clone **(Fig. 1e)**.

**Fig. 1:**
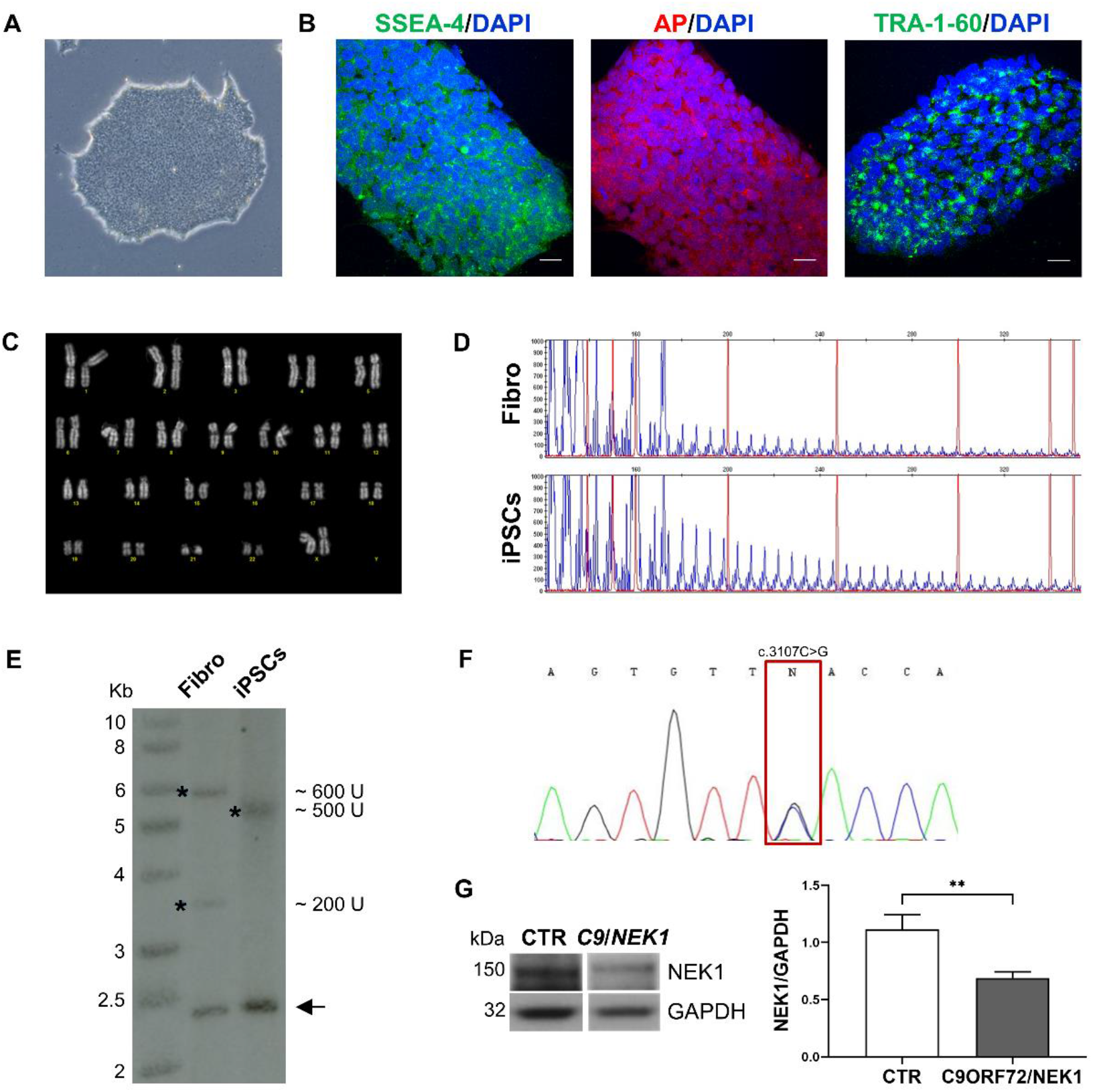
Characterization of the double mutant *C9ORF72*/*NEK1* iPSC line. a) Representative brightfield image of a *C9ORF72*/*NEK1* iPSC colony. b) Representative confocal images of pluripotency markers SSEA-4 (green), Alkaline phosphatase (AP, red) and TRA-1-60 (green) in the *C9ORF72*/*NEK1* iPSC line; nuclei were stained with DAPI (blue). Scale bar = 30 μm. c) Q-banding karyotype of the *C9ORF72*/*NEK1* iPSC line. d) Electropherogram of the *C9ORF72* HRE by Repeat-primed PCR in *C9ORF72*/*NEK1* fibroblasts and iPSCs. e) Southern blot image showing *C9ORF72* HRE in *C9ORF72*/*NEK1* fibroblasts and iPSCs. Asterisks indicate expanded alleles (U, units); arrows indicate the wild-type alleles. f) Electropherogram of Sanger sequencing of *NEK1* gene mutation c.3107C>G:p.Ser1036Ter in the *C9ORF72*/*NEK1* iPSC line. g) Representative Western blot and densitometry analysis of NEK1 protein in the wild-type healthy control (CTR) and in the *C9ORF72*/*NEK1* iPSC lines; all values were normalized on the CTR mean value. GAPDH was used for sample normalization. Mean ±SEM; Student’s t-test (n=3, **p<0.01).

Western Blot analysis showed a significant reduction of NEK1 protein levels (0.6X) in *C9ORF72*/*NEK1* iPSCs compared to an iPSC line derived from a wild-type healthy individual **(Fig. 1g)**, confirming that the *NEK1* p.Ser1036Ter mutation results in NEK1 haploinsufficiency also at protein level.

### 3.2. Double mutant *C9ORF72*/*NEK1* iPSCs show more pathological RNA foci compared to single mutant *C9ORF72* cells

To assess whether the newly generated double mutant *C9ORF72*/*NEK1* iPSC line maintained *C9ORF72*-associated pathological hallmarks, we analysed the presence of the toxic RNA foci generated by the transcription of the *C9ORF72* HRE by RNA-FISH assay. As a positive control for RNA foci formation we used an iPSC line derived from an ALS patient carrying a *C9ORF72* HRE of 1100 units, while a wild-type healthy control iPSC line (CTR) was used as a negative control **(Fig. 2a)**. We found that the *C9ORF72*/*NEK1* iPSC line had a significantly higher percentage of RNA foci-positive cells (58.6%) compared to the *C9ORF72* iPSC line (17.9%) **(Fig. 2b)**, together with a higher mean number of RNA foci per foci-positive cells (9.6 foci in *C9ORF72/NEK1* versus 1.2 foci in *C9ORF72*) **(Fig. 2c)**.

**Fig 2:**
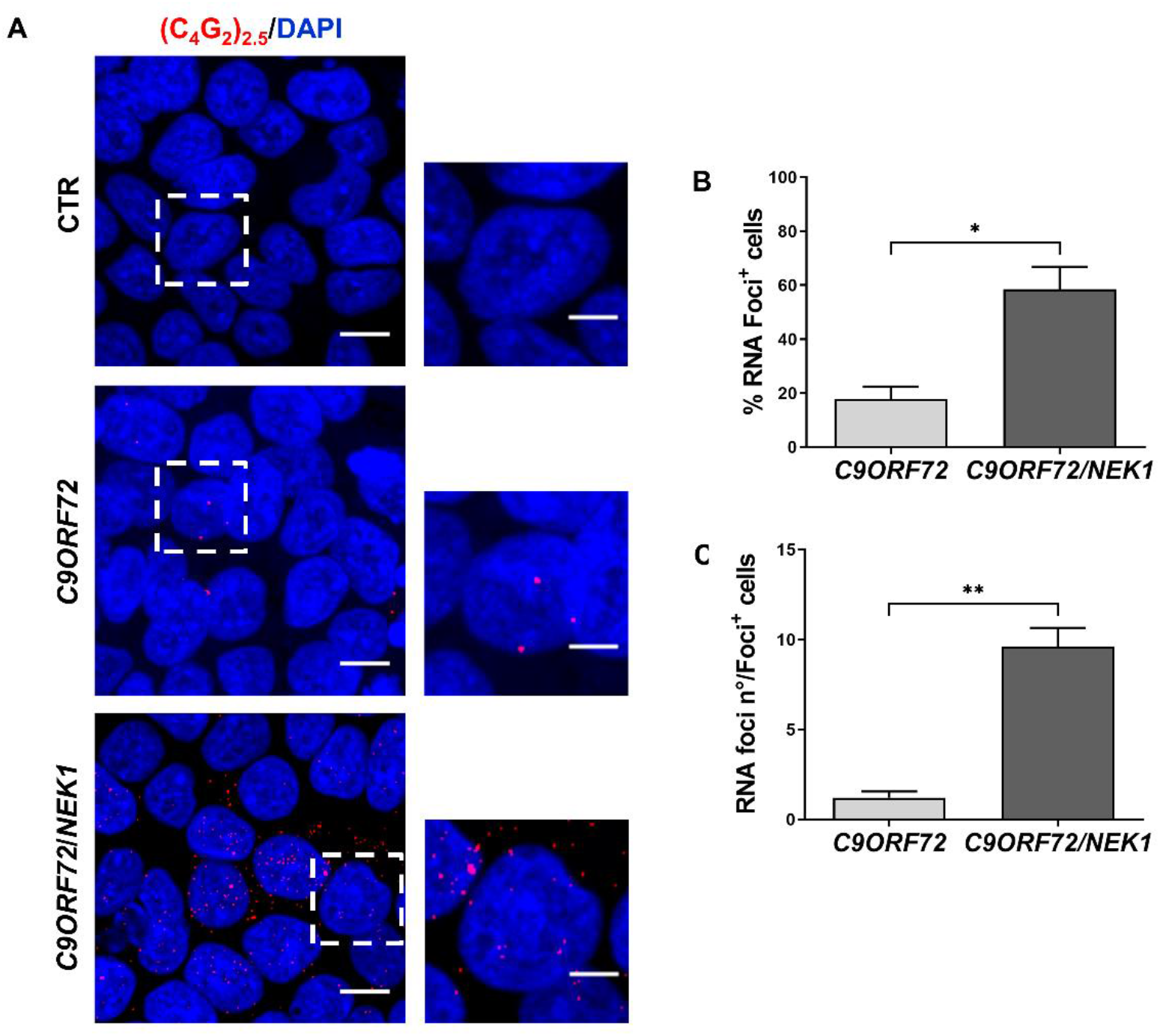
Analysis of *C9ORF72* RNA foci in the double mutant C9ORF72/NEK1 iPSC line. a) Representative images of *C9ORF72* sense RNA foci (red) in wild-type healthy control (CTR), *C9ORF72* and *C9ORF72*/*NEK1* iPSC lines; nuclei were stained with DAPI (blue). Full image (left), scale bar = 10 μm. Inset magnification (right), scale bar = 5 μm. Quantification of b) the percentage of *C9ORF72* RNA foci-positive cells and c) the mean number of *C9ORF72* RNA foci per foci-positive cells in the *C9ORF72* and *C9ORF72*/*NEK1* iPSC lines. Mean ±SEM; Student’s t-test (n=3, >80 cells for each replicate were analysed; *p<0.05, **p<0.01).

### 3.3. Double mutant *C9ORF72*/*NEK1* iPSC-MNs show increased DNA damage levels compared to single mutant *C9ORF72* iPSC-MNs

Given previous literature data about the involvement of mutant *C9ORF72* or *NEK1* genes in the DNA damage pathway (Higelin et al., 2018; Lopez-Gonzalez et al., 2016), we analysed DNA damage response (DDR) in MNs differentiated from the double mutant *C9ORF72*/*NEK1* patient-derived iPSC line. At day 30 of differentiation, all iPSC-MNs from *C9ORF72*/*NEK1, C9ORF72* and wild-type healthy control lines tested positive for the expression of the specific motoneuronal marker choline-acetyltransferase (ChAT) **(Fig. 3a)** and of the neuronal markers βIII-tubulin **(Fig. 3a)** and SMI-312 (pan-axonal neurofilaments) **(Supplementary Fig. S2a)**.

**Fig. 3:**
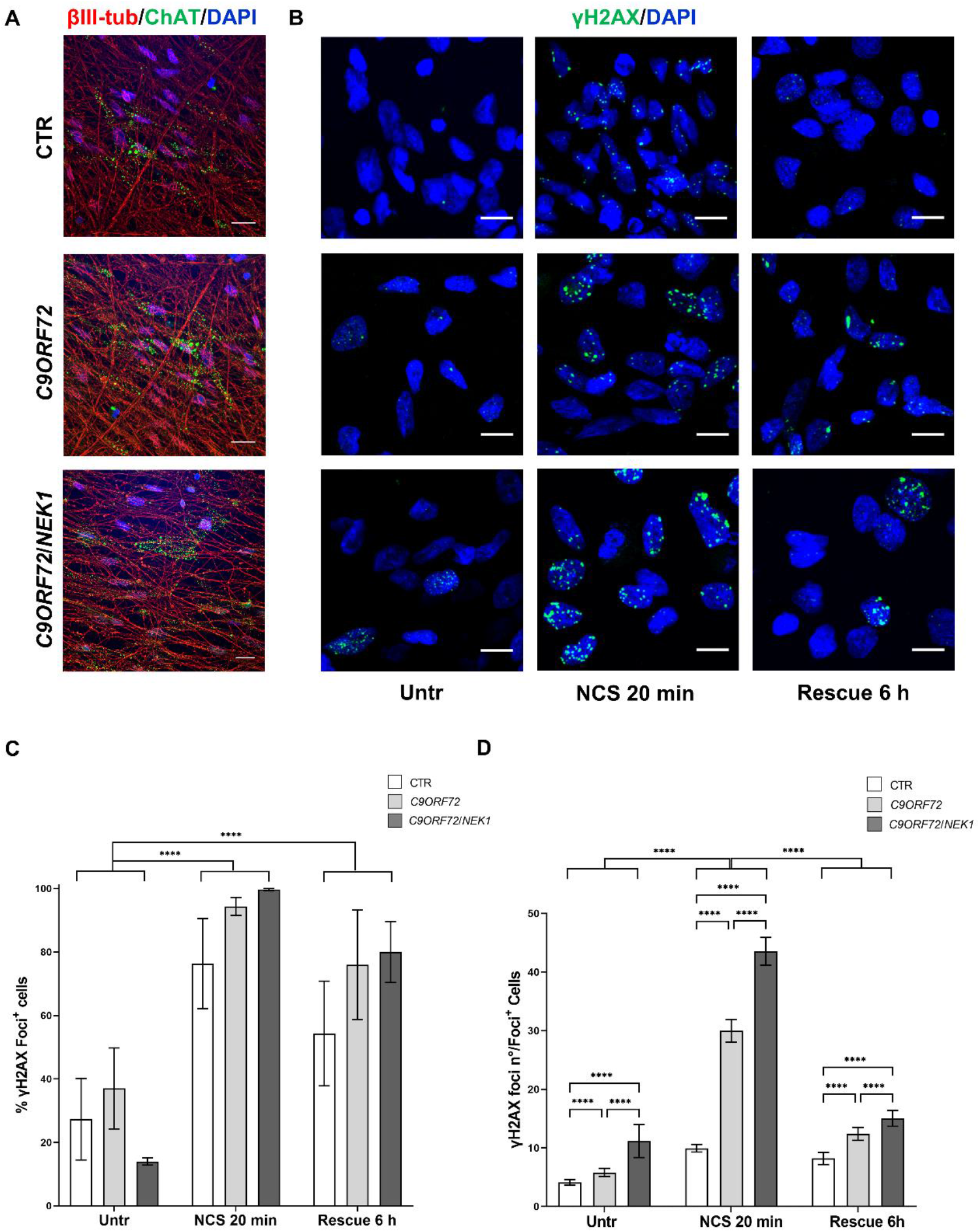
Analysis of DNA damage and DNA damage response (DDR) in *C9ORF72*/*NEK1* iPSC-MNs. a) Representative confocal images of neuronal marker βIII-tubulin (red) and motoneuronal marker ChAT (green) in wild-type healthy control (CTR), *C9ORF72* and *C9ORF72*/*NEK1* iPSC-derived motoneurons (iPSC-MNs); nuclei were stained with DAPI (blue). Scale bar = 20 μm. b) Representative confocal images of γH2AX foci (green) in CTR, *C9ORF72* and *C9ORF72*/*NEK1* iPSC-MNs in untreated conditions (Untr), after Neocarzinostatin treatment for 20 minutes (NCS 20 min) and at 6 hours-rescue time point (Rescue 6 h); nuclei were stained with DAPI (blue). Scale bar = 10 μm. Quantification of c) the percentage γH2AX foci-positive cells and d) the mean number of γH2AX foci per foci-positive cell. Mean ±SEM; Two-way ANOVA, Tukey’s post hoc test (n=3, >100 cells per condition for each replicate were analysed; ****p<0.0001).

To evaluate DNA damage and DDR, we induced single and double strand DNA breaks using the radiomimetic agent Neocarzinostatin (NCS) and investigated DNA damage levels at basal conditions, immediately after 20 minutes treatment with NCS and at 6 hours-rescue time point by staining for the phosphorylated form of the H2AX histone (γH2AX) **(Fig. 3b)**, a well-established DNA damage marker. By quantifying the percentage of γH2AX foci-positive cells, we observed that all iPSC-MN lines had foci-positive cells at basal levels, with no significant differences among the experimental groups (CTR: 27.3%, *C9ORF72*: 37.0%, *C9ORF72*/*NEK1*: 14.0%) **(Fig. 3c)**. After treatment with the NCS agent, the percentage of foci-positive cells significantly increased as expected in all groups (*CTR*: 54.3%, *C9ORF72*: 94.3%, *C9ORF72*/*NEK1*: 99.7%) and remained elevated at 6 hours-rescue (CTR: 54.3%, *C9ORF72*: 76.0%, *C9ORF72*/*NEK1*: 80.0%), without returning to the basal level values **(Fig. 3c)**. Quantification of the mean number of γH2AX foci per foci-positive cells showed that, at basal levels, the *C9ORF72* line had a significantly higher number of foci compared to the CTR line and the *C9ORF72*/*NEK1* iPSC-MNs showed a significant higher number of foci compared to both *C9ORF72* and CTR iPSC-MNs (*CTR*: 4.1 foci, *C9ORF72*: 5.8 foci, *C9ORF72*/*NEK1*: 11.2 foci) **(Fig. 3d)**. After DNA damage induction by NCS, the mean number of foci significantly increased in all groups (CTR: 9.9 foci, *C9ORF72*: 30.0 foci, *C9ORF72*/*NEK1*: 43.6 foci), with the double mutant line showing the greatest increase, and it then significantly decreased at 6 hours-rescue, still maintaining the same differential pattern observed among the three experimental groups (CTR: 8.2 foci, *C9ORF72*: 12.4 foci, *C9ORF72*/*NEK1*: 15.0 foci).

### 3.4. Primary cilia are similarly defective in double *C9ORF72/NEK1* and single *C9ORF72* mutant iPSC-MNs

Since recessive *NEK1* gene mutations cause human skeletal ciliopathies (Thiel et al., 2011) and NEK1 protein is involved in regulating ciliogenesis (Shalom et al., 2008; White and Quarmby, 2008), we decided to investigate primary cilia formation in the double mutant iPSC-MNs in comparison to *C9ORF72* and CTR iPSC-MNs. Primary cilium was visualized by IF with adenylate cyclase III (AC-III) marker **(Fig. 4a)**. By quantifying the percentage of cilia-positive cells, we found that both single mutant *C9ORF72* and double mutant *C9ORF72/NEK1* iPSC-MNs showed a significant decreased percentage of cilia-positive cells compared to CTR iPSC-MNs (CTR: 52.4%, *C9ORF72*: 26.7%, *C9ORF72*/*NEK1*: 25.9%) **(Fig. 4b)**. Image analysis of cilium size revealed that *C9ORF72* iPSC-MNs showed a significant shorter cilium length compared to CTR iPSC-MNs (CTR: 1.5 μm, *C9ORF72*: 1.0 μm) and that also *C9ORF72*/*NEK1* iPSC-MNs presented significantly shorter cilia (0.9 μm) **(Fig. 4c)**. However, we found no significant differences between the single mutant *C9ORF72* and the double mutant *C9ORF72/NEK1* iPSC-MNs, neither in the percentage of cilia-positive cells **(Fig. 4b)**, nor in the cilium length **(Fig. 4c)**.

**Fig. 4:**
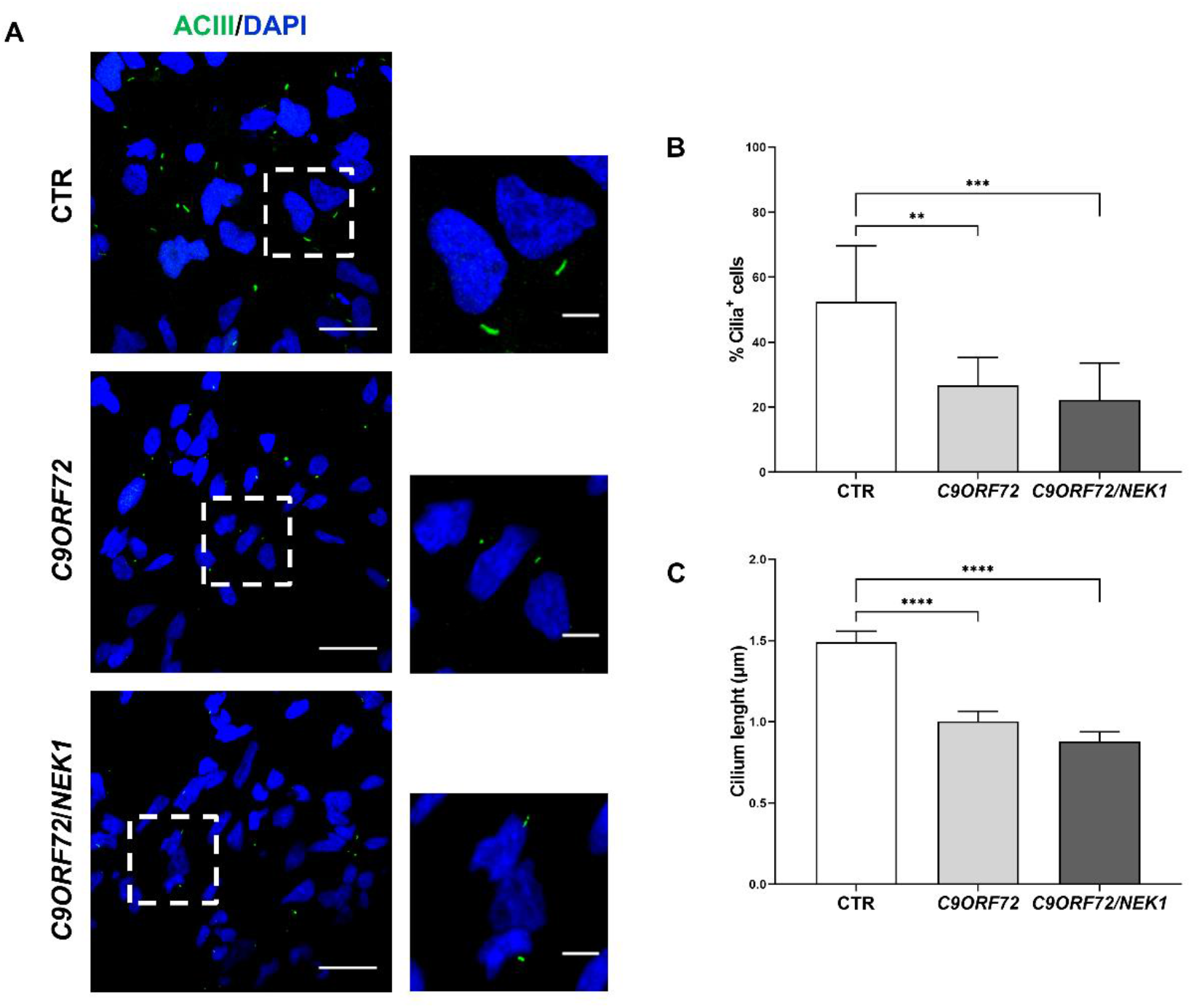
Analysis of primary cilia in *C9ORF72* and *C9ORF72*/*NEK1* iPSC-MNs. a) Representative confocal images of primary cilium marker AC-III (green) in wild-type healthy control (CTR), *C9ORF72* and *C9ORF72*/*NEK1* iPSC-motoneurons (iPSC-MNs); nuclei were stained with DAPI (blue). Full image (left), scale bar = 10 μm; inset magnification (right), scale bar = 5 μm. Quantification of b) the percentage of primary cilia-positive cells and c) the mean cilium length. Mean ±SEM; One-way ANOVA, Tukey’s post hoc test. (n=3, >200 cells for each replicate were analysed; **p<0.01, ***p<0.001, ****p<0.0001).

## 4. Discussion

In this study we analysed the biological effects of *NEK1* haploinsufficiency as a possible modifier of *C9ORF72* pathology by using *in vitro* disease models such as iPSCs obtained from an ALS patient carrying a concomitant *C9ORF72* HRE and a rare *NEK1* LOF variant. The nonsense mutation p.Ser1036Ter in *NEK1* gene has already been described to be associated to ALS aetiology in cohorts of different origins (Ruf et al., 2023) and also in combination with *C9ORF72* HRE in a patient with familiar ALS, leading to a near complete loss of the mutated *NEK1* transcript, likely via nonsense-mediated mRNA decay (Nguyen et al., 2018b). However, there is still no functional evidence if and how this variant acts as a possible modifier on an already *C9ORF72*-mutated genetic background.

In the iPSC line generated from the double mutant ALS patient we confirmed *NEK1* haploinsufficiency, further confirming a LOF mechanism for this variant also at protein level. Interestingly, the double mutant *C9ORF72*/*NEK1* iPSC line showed an increased number of cells forming pathological *C9ORF72*-associated RNA foci together with a higher number of RNA foci compared to the single mutant *C9ORF72* iPSC line. There is no clear evidence in literature linking *NEK1* gene to an altered RNA metabolism that could favour the formation of pathological RNA foci. However, *NEK1* is involved in DNA damage response and *C9ORF72* RNA foci and pathological DPRs are reported to induce double strand DNA breaks and R-loops (DNA:RNA hybrids) formation (Walker et al., 2017), so that a link between RNA foci transcription and NEK1 haploinsufficiency may be plausible, although still to be demonstrated mechanistically.

In line with this view, the analysis of DNA damage at baseline showed increased levels in the *C9ORF72*/*NEK1* double mutant cells compared to *C9ORF72* single mutant iPSC-MNs, although the response to DNA damage by NCS agent was not affected in either of the two mutant lines. However, a previous study reported defective DDR in both *C9ORF72* and *NEK1* LOF (p.Arg812Ter) single mutant iPSC-MNs (Higelin et al., 2018). This discrepancy could be due to the use of different genotoxic agents (NCS versus γ-irradiation) or to a different susceptibility of the obtained iPSC-MNs to DNA damage induction, which may also depend on the distinct protocols used to differentiate MNs in the two studies.

We here report for the first time that the *C9ORF72* HRE affects primary cilia formation in ALS-patient derived iPSC-MNs. In neuronal cells the primary cilium acts an antenna-like organelle responding to external stimuli and regulating several cell pathways, including autophagy (Mill et al., 2023). Impairment of ciliogenesis and ciliary signalling has been recently associated with neurodegenerative disorders, including Parkinson’s and Alzheimer’s diseases (Ma et al., 2022) while, in ALS, the presence of defective cilia was described only in transgenic SOD1 mice so far (Ma et al., 2011).

Our findings supports the recent observation that C9ORF72 protein, in complex with SMCR8, acts as a major negative regulator of primary ciliogenesis (Tang et al., 2023). Nonetheless, our data indicate that patient-derived *C9ORF72* iPSC-MNs show shorter cilium length and not longer ones, as recently described in cell lines or mice knocked-down for *C9ORF72* gene (Tang et al., 2024, 2023) suggesting that not only *C9ORF72* haploinsufficiency, but also RNA and DPRs gain-of-function mechanisms may influence ciliogenesis regulation. Our results also show that the presence of an additional *NEK1* LOF mutation does not worsen *C9ORF72* negative effect on primary cilia formation, suggesting a not additive effect of NEK1 haploinsufficiency on ciliogenesis in contrast to what observed for DNA damage. However, preliminary data from our group report that *NEK1* haploinsufficiency alone negatively impairs ciliogenesis in iPSC-MNs and brain organoids (https://doi.org/10.1101/2024.02.29.582696), suggesting that the interplay between mutant *NEK1* and *C9ORF72* genes in these two cell pathways needs further investigation. Recently, it has been also shown that *NEK1* haploinsufficiency disrupts microtubule dynamics (Mann et al., 2023), further supporting a role of NEK1 protein in directly or indirectly regulating primary cilium formation and length by binding to microtubules.

Despite having identified a biological impact of the *NEK1* LOF mutation on different cellular pathways in *in vitro* patient-derived *C9ORF72* disease models, including increased pathological *C9ORF72* RNA foci formation and DNA damage, we did not observe any specific clinical feature which could distinguish the double mutant *C9ORF72*/*NEK1* ALS patient from those harbouring only *C9ORF72* HRE. This certainly sustains that ALS disease aetiology is also very complex in familial or mutation-associated cases with a variety of still-to-be-identified modifiers, including additional genetic risk factors and epigenetic modifications, which may determine the final clinical outcome and account for the vast heterogeneity observed among ALS patients. Nevertheless, our study supports the use of patient-derived iPSCs as suitable *in vitro* disease models to investigate the contribution of multiple genetic variants as possible modifying factors in *C9ORF72*-associated pathology with the aim to functionally assess their biological relevance in the context of an oligogenic condition.

## Declarations

## Acknowledgments

SS, SI and VC were recipients of fellowships from the PhD program in “Experimental Medicine”, Università degli Studi di Milano. The authors acknowledge Dr. Claudia Fallini (University of Rhode Island, USA) and Dr. Sofia Francia (IGM-CNR, Italy) for their helpful suggestions and Dr. Jan Hensen (University of Bonn, Germany) for his help with the CiliaQ plugin.

AR acknowledges “Aldo Ravelli Center for Neurotechnology and Experimental Brain Therapeutics”, Università degli Studi di Milano; VS acknowledges the ERN Euro-NMD; NT acknowledges PSR 2021 grant, Università degli Studi di Milano, for financial support.

## Funding

The project has been financially supported by the Italian Ministry of Health (GR-2016-02364373). The publication fee has been supported by Ricerca Corrente from the Italian Ministry of Health.

## Competing interests

The authors declare no competing interests.

## Authors’ contributions

Conceptualization: AR, SS, SI. Investigation and methodology: SS, SI, MNS, VC, SP, CC, NT and PB. Formal analysis: SS, SI and AB. Writing-original draft: AR, SS and SI. Supervision: AR and VS. Writing-review & editing: all authors.

## Availability of data and material

Experimental raw data are available on Zenodo (DOI 10.5281/zenodo.10650632).

## Supplementary material

**Supplementary Table S1.** List of primary and secondary antibodies for WB and IF analyses.

| **WB** | **Supplier** | **Dilution** |
| --- | --- | --- |
| GAPDH | Santa Cruz Biotechnology (sc-47724) | 1:500 |
| NEK1 | Santa Cruz Biotechnology (sc-398813) | 1:200 |
| HRP-conjugated anti-mouse | Sigma-Aldrich (A9309) | 1:20'000 |
| **IF** | **Supplier** | **Dilution** |
| Adenylate cyclase III | Santa Cruz Biotechnology (sc-518057) | 1:100 |
| Alkaline Phosphatase | Abcam (ab108337) | 1:250 |
| Beta III-Tubulin | Abcam (ab52623) | 1:500 |
| Choline-Acetyltransferase | Chemicon (MAB305) | 1:200 |
| Phospho-Histone H2A.X (Ser139) | Millipore (05-636) | 1:800 |
| SMI-312 | Covance (SIG-32248) | 1:5000 |
| SSEA4 | Invitrogen (14-8843-80) | 1:100 |
| TRA-1-60 | Invitrogen (14-8863-80) | 1:125 |
| Alexa Fluor 488 anti-mouse | Invitrogen (A11034) | 1:500 |
| Alexa Fluor 555 anti-rabbit | Thermo Fisher Scientific (A21430) | 1:500 |

**Supplementary Table S2.** Clinical features of the double mutant *C9ORF72/NEK1* ALS patient.

| **Gender** | **Age at onset** | **Site of onset** | **Cognitive status** | **Family history** | **Survival (months)** |
| --- | --- | --- | --- | --- | --- |
| Female | 50 | Bulbar | ALSbi* | Mother with motoneuron disease, sister with multiple sclerosis | 27 |

\*ALS with behavioral impairment, based on the performance at ECAS and according to the Strong criteria (Strong et al., 2017).

**Supplementary Fig. S1.**
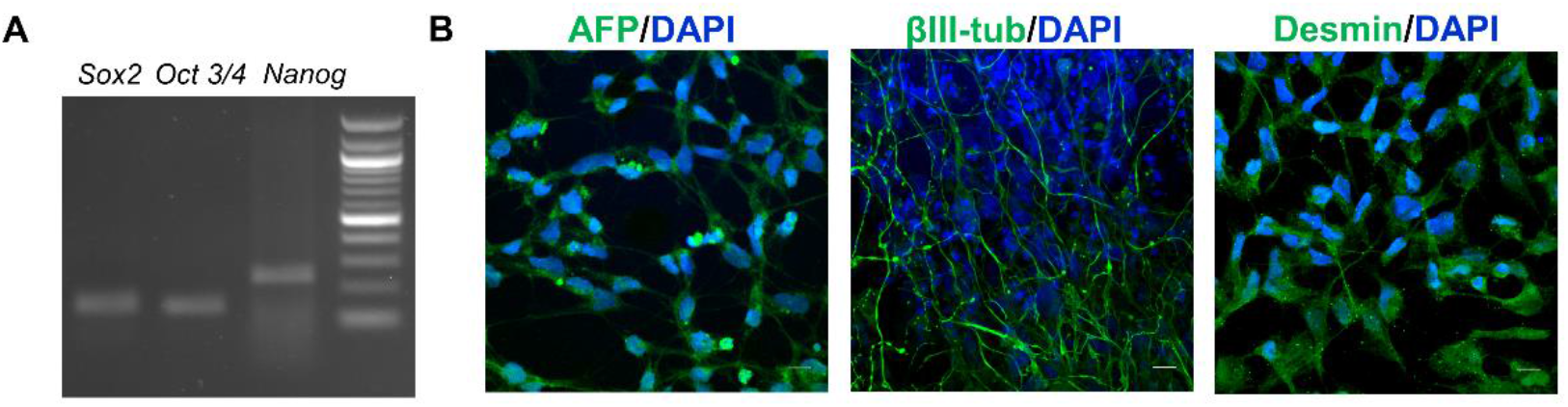
Characterization of the double mutant *C9ORF72*/*NEK1* iPSC line. a) iPSC colonies expressing *SOX2, OCT 3/4* and *NANOG* markers by RT-PCR. b) Representative confocal images of cells expressing the three germ layers markers (all in green): alpha-fetoprotein (AFP), βIII-tubulin (βIII-tub) and desmin in the *C9ORF72*/*NEK1* iPSC line; cell nuclei were stained with DAPI (blue). Scale bar = 30 μm.

**Supplementary Fig. S2.**
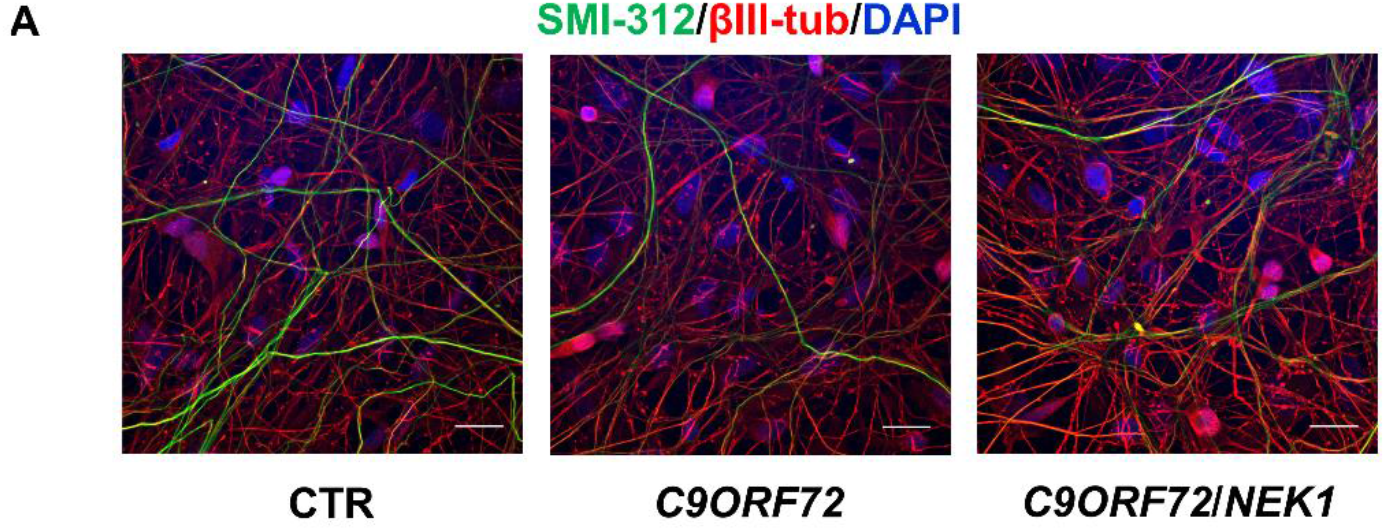
Characterization of the differentiated *C9ORF72*/*NEK1* iPSC-MNs. a) Representative confocal images of neuronal markers SMI-312 (green) and βIII-tub (red) in wild-type control (CTR), *C9ORF72* and *C9ORF72*/*NEK1* iPSC-MNs at day 30 after embryoid bodies dissociation; cell nuclei were stained with DAPI. Scale bar = 20 μm.

## Notes

### Competing Interest Statement

The authors have declared no competing interest.

